# C/EBPα acts as an RNA binding protein essential for its chromatin recruitment and promotes Macrophage differentiation

**DOI:** 10.1101/2021.10.20.465166

**Authors:** Yanzhou Zhang, Mahmoud Bassal, Daniel Friedrich, Simone Ummarino, Tom Verbiest, Junyan Zhang, Danielle Tenen, Arthanari Haribabu, Daniel G Tenen

## Abstract

C/EBPα has known to be a transcription factor that involved in Neutrophil differentiation for decades. However, exploring the Chromatin RNA Immunoprecipitation Sequencing (RIP), we discover that C/EBPα is a RNA binding protein mainly interacts with RNA introns. Structure study and RNA electrophoretic mobility shift assay (REMSA) show that C/EBPα interacts with RNA through two novel RNA binding domains distinct from its DNA binding domain. Mouse bone marrow transplantation and *in vitro* cytokine assay reveal that C/EBPα RNA binding is critical for Macrophage differentiation but not Neutrophil differentiation. Mechanically, RNA binding domains control specific gene transcription. In particular, PU.1 intron 4 RNA interacts with C/EBPα and recruit C/EBPα to its enhancer site, which facilitate PU.1 expression. Taken together, C/EBPα is demonstrated to be a RNA binding protein with unique function distinct from its DNA binding activity. Our finding transforms our knowledge of transcriptional regulation by transcription factor.

**One-Sentence Summary:** C/EBPα interacts with RNA through novel RNA binding domains and regulates PU.1 expression during Macrophage differentiation.

## Introduction

Recent studies on non-canonical RNA binding proteins (RBP) has revealed the RNA binding activity in traditional DNA binding proteins (Di Ruscio et al., 2013; Holmes et al., 2020; Saldana-Meyer et al., 2019; Sigova et al., 2015; Wai et al., 2016; Xu et al., 2021). They may interacts with RNA and DNA similarly or differently, and their RNA binding activity may support their role or provide different functions. For example, DNA methyltransferase 1 (DNMT1) interacts with non-coding RNA *ecC/EBPα* which inhibits its catalytic activity and transcriptional control (Di Ruscio et al., 2013). Transcription factor Yin-Yang 1 (YY1) interacts with RNA transcripts that enhances its occupancy on chromatin elements (Sigova et al., 2015; Wai et al., 2016). CCTTC-binding factor (CTCF) interacts with RNA through zinc finger domain and promotes long term chromatin interactions (Saldana-Meyer et al., 2019). All of these examples highlight the pleiotropic potential of RNA binding activity in regulating distinct functions of traditional DNA binding proteins.

CCAAT-enhancer binding protein alpha (C/EBPα) is a canonical transcription factor which plays a non-redundant role in the transition from common myeloid progenitors (CMPs) to granulocyte macrophage progenitors (GMPs) (Zhang et al., 2004). Knockout C/EBPα in mice abolishes GMPs and results in a complete block of neutrophilic development (Zhang et al., 2004). In the past few decades, the comprehensive picture of C/EBPα function, predominantly focus on its DNA binding activity has merged. A crystal structure of C/EBPα C-terminal bZIP domain DNA binding complex has been shown (Miller et al., 2003; Vinson et al., 1989), and mutation on bZIP domain disrupts DNA binding and therefore disrupts C/EBPα function on inhibiting E2F1 and on inducing neutrophil differentiation (D’Alo et al., 2003; Porse et al., 2001). However, the study on C/EBPα N-terminal region is still limited. Additionally, previous study has shown that bZIP containing protein GCN4 and c-Jun is capable to bind to RNA (Nikolaev et al., 2010; Nikolaev and Pervushin, 2012), which drives a potential capability of C/EBPα RNA association. Taken together, the function of C/EBPα N-terminal region, and its capability and functionality in RNA binding is still remain unclear.

In this study, we surprisingly found that C/EBPα is an RBP which interacts with intron of RNA transcript. Employing alphaFold prediction and RNA electrophoretic mobility shift assay (REMSA), we identified two novel RNA binding domains located in N terminal region distinct from bZIP domain. Abolishing C/EBPα RNA binding activity affected macrophage differentiation but not neutrophil differentiation in bone marrow transplantation assay and in vitro cytokine assay. Mechanically, abolishing C/EBPα RNA binding activity decreased the recruitment of C/EBPα to PU.1 enhancer site, and downregulated PU.1 expression. Correspondingly, macrophage related genes such CD163, CD206, iNOS, IL1β, TNFα, IL10, Arg-1 and CCL-2 were also decreased. In the past few decades, C/EBPα is known as a major regulator for neutrophil (Ma et al., 2014; Zhang et al., 1997; Zhang et al., 1998; Zhang et al., 2004). The role of C/EBPα in macrophage differentiation is remain unclear. Our study shows C/EBPα is a non-canonical RBP involves in macrophage differentiation which transforms our knowledge underlying hematopoiesis.

## Results

### C/BEPα is a RNA binding protein associated with RNA introns

To investigate whether transcription factor C/EBPα also function as a RNA binding protein, we first performed C/EBPα chromatin RNA immunoprecipitation followed by RNA sequencing (RIP-seq) (Fig. 1a) in both HL60 cells (Fig. 1b) and THP-1 cells (Fig. 1c). The results show that C/EBPα mainly interacts with introns of coding RNA. To validate the sequencing results, we first validated the specificity of C/EBPα antibody and ability to specifically immunoprecipate C/EBPα with magnetic beads using western blot (fig. S1d). We next performed chromatin RNA immunoprecipitation followed by quantitative real time PCR (RIP-qRT-PCR) and UV-crosslinked RNA immunoprecipitation followed by quantitative real time PCR (CLIP-qRT-PCR). We first observed UV-crosslinked C/EBPα RNA complexes migrated in a less defined band, indicating crosslinked with heterogeneous RNAs (fig. S1a). PU.1, CTBP1, and NEAT1 were then selected as their peaks are significant higher compare to IgG control and Input (Fig. 1d). Enrichment of C/EBPα on positive binding region but not on negative binding region were observed in both RIP-qRT-PCR (Fig. 1e) and CLIP-qRT-PCR (fig. S1b-c). Interestingly, we identified three highly similar CA-rich motif, which occupied large percent of RIP-seq peaks (Fig. 1f). Importantly, these motifs were highly repeated in binding regions of targeted introns but not in other part of the introns (fig. S1e-f).

**Figure. 1.**
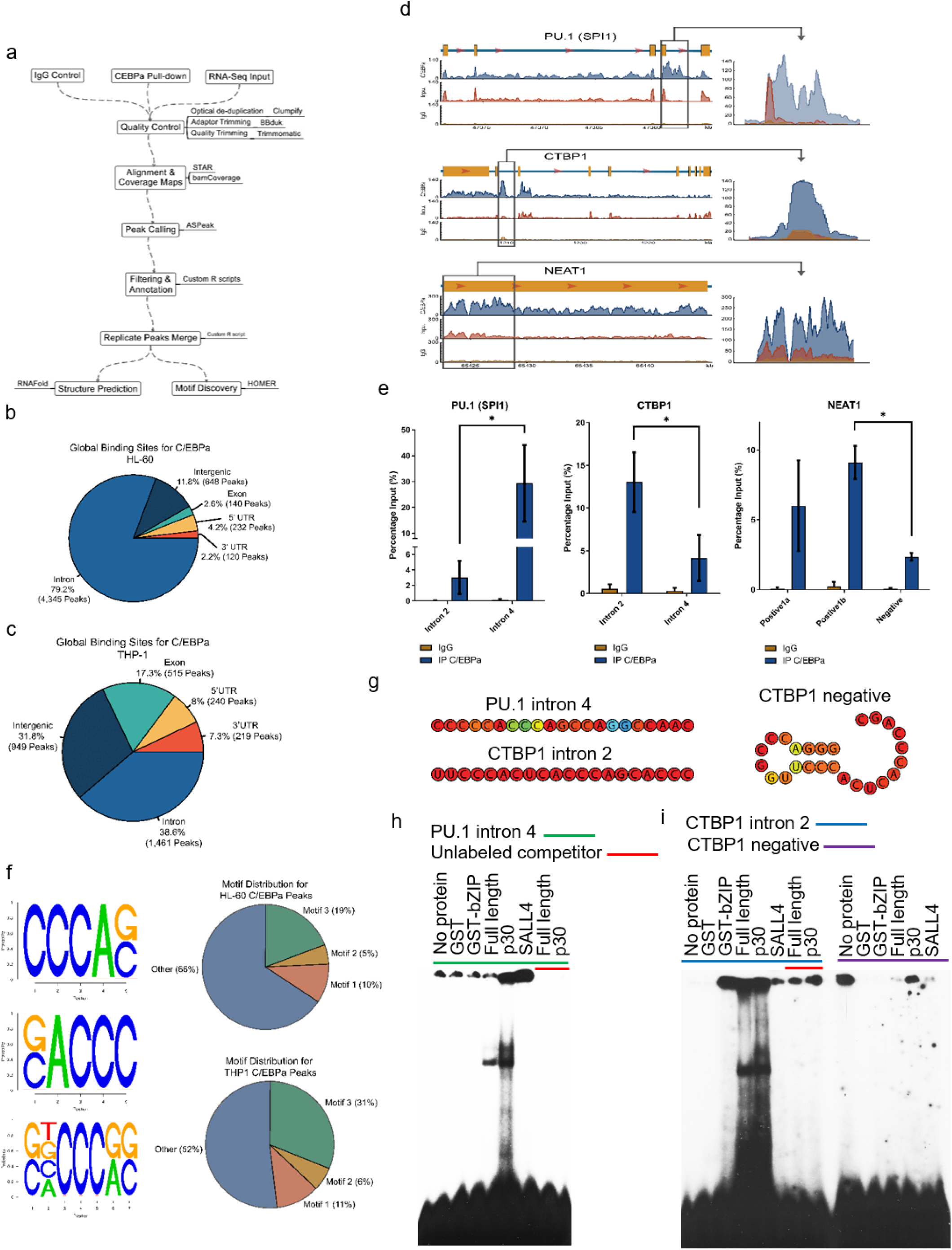
C/EBPα binds intron of RNA transcript directly. (a) Pipeline of chromatin RNA immunoprecipitation (RIP) analysis. (b, c) Pie chart showing C/EBPα mainly interacts with RNA introns in both HL60 cells (b) and THP-1 cells (c). (d) Representative peaks of C/EBPα RNA binding on PU.1 intron 4 (upper), CTBP1 intron 2 (middle) and NEAT1 (down) viewed on IGV. (e) RIP-qRT-PCR validate of C/EBPα enrichment on PU.1 intron 4 (left), CTBP1 intron 2 (middle) and NEAT1 (right) RNAs. C/EBPα enrichment was shown on positive region but no on negative region. (f) Pie chart showing the CA-rich binding motif (left) occupied large percent of C/EBPα RIP peaks in both HL60 cells (upper) and THP-1 cells (down). (g) Sequence and predict structure of RNA probes used in REMSA. (h, i) REMSA blots show both C/EBPα p42 and p30 isoforms interact with PU.1 intron 4 RNA (h) and CTBP1 intron 2 RNA (i) directly *in vitro*, but not interacts with CTBP1 negative probe (i).

We further confirmed that C/EBPα is a RNA binding protein by observing direct RNA interaction between C/EBPα and RNA *in vitro*. We selected three small RNA probes from PU.1 intron 4 (Fig. 1g and fig. S1e) and CTBP1 intron 2 (Fig. 1g and fig. S1f) based on the CA-rich motif, and performed RNA electrophoretic mobility shift assay (REMSA) with bacteria purified recombinant C/EBPα proteins. We observed signal shift when linear RNA probes containing multiple repeat of CA-rich motif mixed with either C/EBPα p42 or p30 variants (Fig. 1h-i), and the shift signals disappear when unlabeled competitor was added to the mixture (Fig. 1h-i). Most importantly, there was no shift signal been observed when proteins was mixed with loop structured probe (Fig. 1i), suggesting that the CA-rich motif was more accessible in linear form of RNA. These data not only confirmed C/EBPα directly interacts with RNA but also suggest that C/EBPα may interacts with specific RNA introns during cellular process.

### C/EBPα interacts with RNA through N terminal region but not bZIP domain

We next investigated which domain of C/EBPα is responsible for RNA binding. We first performed REMSA experiment by mixing RNA probes with different deletion mutation of C/EBPα (Fig. 2a). Unexpectedly, we observed shift signal in almost all the deletion mutations of C/EBPα except bZIP domain fragment (Fig. 2b-c), suggesting that bZIP domain of C/EBPα is not responsible for RNA binding. However, bZIP domain binds to neutrophil elastase promoter DNA very well (fig. S2a), indicating that the purified bZIP domain is functioning well. Unlike other transcription factor such as SOX2 or c-Jun which interacts RNA and DNA through same domain (Holmes et al., 2020; Nikolaev et al., 2010), C/EBPα tend to separate the RNA binding and DNA binding activity. To further confirm, we generated a C/EBPα basic domain mutation (BRM2) construct (Fig. 2a) which proved to reduce the DNA binding ability (Kowenz-Leutz et al., 2016). However we observed strong RNA binding activity in BRM2 mutation, further confirming that DNA binding activity and RNA binding activity of are separated in C/EBPα. As a control for REMSA, we also designed a DNA probe with the same sequence as the RNA probe, we observed minimum shift signal in DNA compare to RNA (Fig. 2b-c), suggesting that C/EBPα interacts with RNA specifically.

**Figure. 2.**
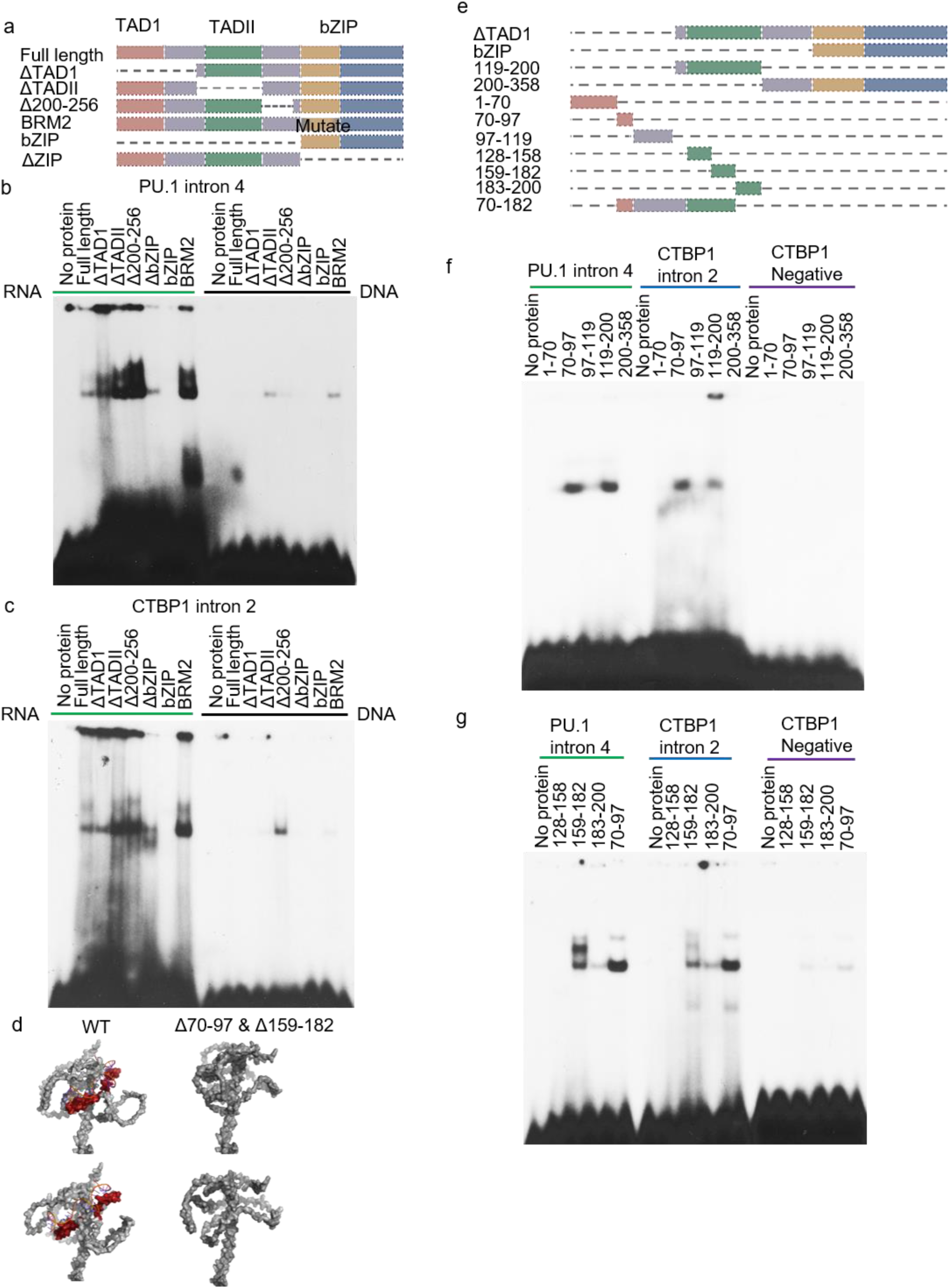
C/EBPα interacts with RNA through two novel RNA binding domain located in N-terminal region but not through bZIP domain. (a) Schematic shows C/EBPα single domain deletions. (b, c) REMSA blot show C/EBPα bZIP domain is not responsible for RNA binding of PU.1 intron 4 (b) and CTBP1 intron 2 (c). DNA probes contains exactly same sequence of RNA probe, no binding was observed on these DNA probes indicating the specificity of C/EBPα RNA binding. (d) AlphaFold prediction of the structure of C/EBPα-RNA binding complex indicate the two novel RNA binding domain (aa. 70-97 and aa. 150-182). Left: C/EBPα wild type which shows a unique RNA pocket. Right: C/EBPα RNA binding domain deletion which shows close and collapse of RNA pocket. (e) Schematic shows C/EBPα fragments. (f, g) REMSA shows C/EBPα aa. 70-97 and aa. 159-182 interacts with RNA directly *in vitro*.

To further narrow down the RNA binding domain of C/EBPa, we performed a nuclear magnetic resonance (NMR) with C/EBPα N terminal region and RNA probe used in REMSA. Compare to C/EBPα along, we observed chemical shift when C/EBPα protein mixed with RNA (fig. S2b), confirming C/EBPα interacts with RNA directly *in vitro*. Unfortunately, NMR result show that C/EBPα N terminal is too dis-ordered in NMR buffer to assign the binding domain. However, we still observed RNA binding activity, suggesting that C/EBPα RNA binding activity may not require a highly stable binding structure like its DNA binding activity (Miller et al., 2003).

We next performed AlphaFold prediction for C/EBPα RNA binding (Jumper et al., 2021). Interestingly, although the N terminal of C/EBPα is dis-ordered, we observed a RNA pocket formed within its N terminal region that was able to capture linear RNA (Fig. 2d). Based on the prediction, there were two small regions that was able to interact with RNA and was very important to maintain this RNA pocket (Fig. 2d). To validate the prediction, we purified several C/EBPα fragment (Fig. 2e) and performed REMSA experiment. Interestingly, we observed shift signal in two predicted fragments which are aa. 70-97 and aa. 159-182 but not in other fragments (Fig. 2f-g), indicating that C/EBPα interacts RNA through these two RNA binding domains. To further confirm, we overexpressed C/EBPα wild type or C/EBPα double RNA binding domain deletion mutation into HL-60 cells and performed RIP-qRT-PCR (fig. S2c), we observed that the RNA binding activity is disappear in C/EBPα double RNA binding domain deletion mutation. The pull down efficiency between wild type and mutation proteins were examined by western blot (fig. S2d). We next performed AlphaFold prediction again with C/EBPα double RNA binding domain deletion mutation, interestingly we observed collapse and closure of the RNA pocket (Fig. 2d). These data demonstrate two novel RNA binding domains localized in C/EBPα N terminal region.

### C/EBPα interacts with RNA and DNA with similar affinity

Since C/EBPα interacts with RNA through novel RNA binding domains distinct from DNA binding, we were interested in measuring the affinity of this RNA binding. Isothermal titration calorimetry (ITC) was performed using C/EBPα RNA binding domains (Fig. 3a), we calculated *Kd* of RNA binding domain 1 (aa. 70-97) is around 11.5 µM and *Kd* of RNA binding domain 2 (aa. 159-182) is around 16 µM. The N terminal fragment that contains both RNA binding domains has the highest affinity with *Kd* around 5.8 µM. As a control, the non-RNA-binding fragments (aa. 1-70 and bZIP domain) did not show any binding activity on ITC (fig. S2e).

**Figure. 3.**
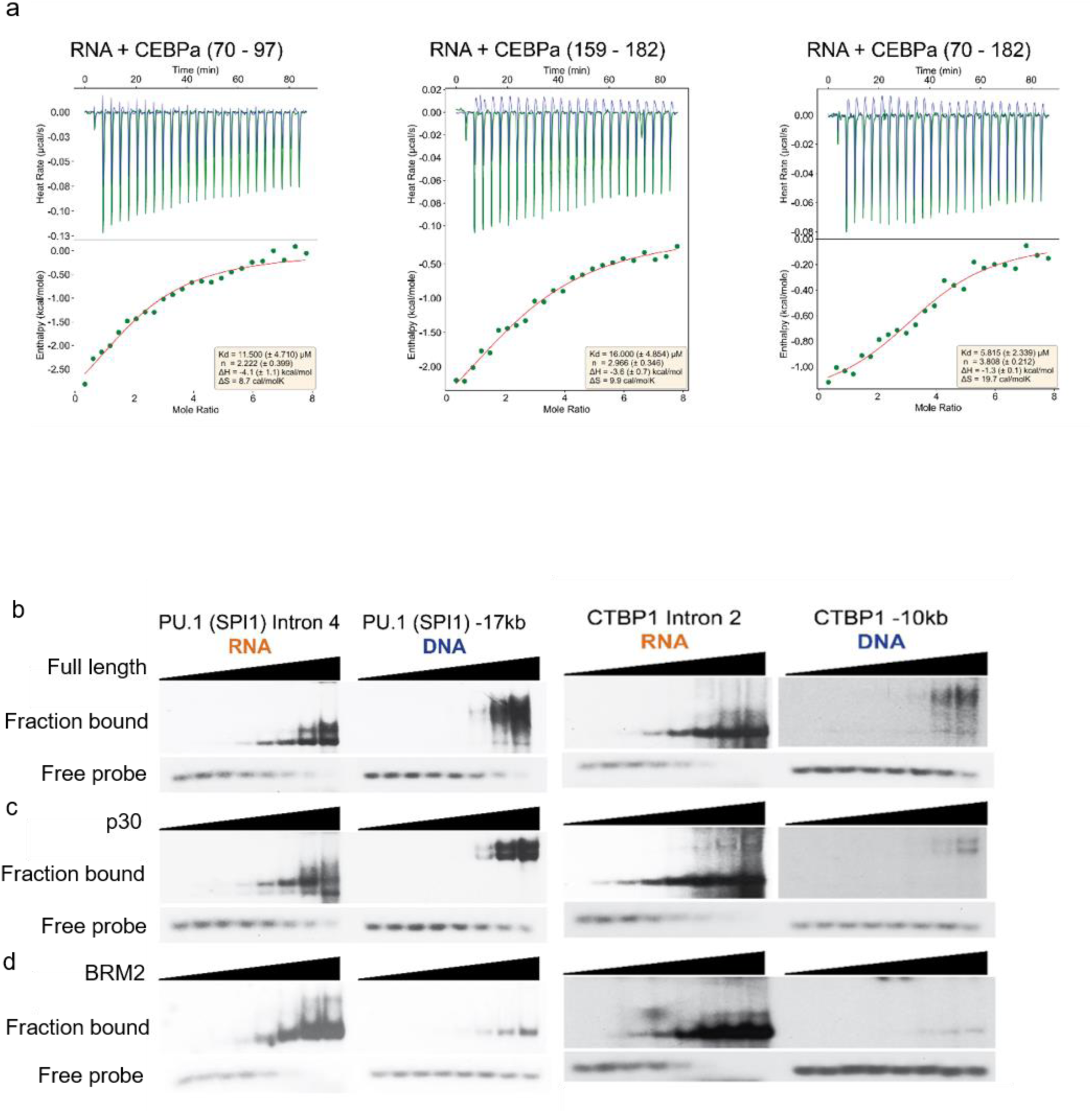
C/EBPα binds RNA and DNA with similar affinity. (a) Isothermal titration calorimetry (ITC) affinity calculation of C/EBPa RNA binding domain binds to RNA. aa. 70-97 *Kd* 11.5 µM (left), aa. 159-182 *Kd* 16 µM (middle), aa. 70-182 *Kd* 5.8 µM (right). (b, c) REMSA blot shows similar binding affinity of C/EBPα full length (b) and p30 (c) between RNA and DNA. (d) REMSA blot shows C/EBPα BRM2 mutation reduce DNA binding but has no effect on RNA binding.

Since C/EBPα is able to bind both RNA and DNA, we compared the binding affinity between them by performing REMSA experiment. With increased amount of C/EBPα p42 (Fig. 3b) and C/EBPα p30 (Fig. 3c), we observed similar increase of shift signal and similar decrease of free probe signal, indicating that C/EBPα binds RNA and DNA with similar affinity. Additionally, C/EBPα BRM2 mutation (Fig. 3d) binds RNA with similar affinity with C/EBPα wild type variants, whereas its DNA binding affinity is much lower than wild type variants, further demonstrating that C/EBPα RNA binding activity and DNA binding activity are separate.

### C/EBPα RNA binding activity contribute to Macrophage differentiation but not Neutrophil differentiation

We next investigated whether C/EBPα RNA binding activity is essential for hematopoiesis. We performed mouse bone marrow transplantation experiment. Lin^-^Sca-1^+^c-kit^+^ cells (LSKs) isolated from poly (inosinic acid) poly (cytidylic acid) (poly I:C) treated Mx.1-Cre^+^C/EBPα^f/f^ C/EBPα conditional knockout mice (KO) were transduced with MigR1-vector (vector), MigR1-C/EBPα-wild type-ER-IRES-GFP (WT), MigR1-C/EBPα-Δ70-97+Δ159-182-ER-IRES-GFP (Δ70-97+Δ159-182), MigR1-C/EBPα-Δ70-97-ER-IRES-GFP (Δ70-97) or MigR1-C/EBPα-ΔTADII-ER-IRES-GFP (ΔTADII) retrovirus, and transplanted into lethally irradiated congenic recipients along with a radio-protective dose of congenic mice lineage depleted bone marrow cells (Fig. 4a). After reconstitution of the congenic blood system (fig. S3a), tamoxifen was feed to transplanted mice induce C/EBPα-ER translocation from cytoplasma to nuclear (Truong et al., 2003; Umek et al., 1991). Effect of C/EBPα on hematopoiesis was monitored by flow cytometry of spleen and bone marrow (fig. S3b). Knockout of C/EBPα (KO) show complete loss of neutrophil population (CD11b^high^Ly6G^+^) (fig. S3b). Except vector control (n=6), myeloid cells population (CD11b^intermediate-high^B220^-^CD4^-^CD8^-^) were successfully rescued in spleen (fig. S4) and bone marrow (fig. S5) in both C/EBPα wild type (n=8) and deletion mutations. Additionally, RNA binding domain deletion mutation of C/EBPα (Δ70-97+Δ159-182) (n=6) was able to rescue neutrophil population at a similar level as C/EBPα wild type (fig. S6), suggesting that disrupting RNA binding activity has minor effect on neutrophil differentiation. Whereas disrupting DNA binding activity has a major effect on neutrophil differentiation, indicating that the function of C/EBPα RNA binding activity may distinct from its DNA binding activity.

**Figure. 4.**
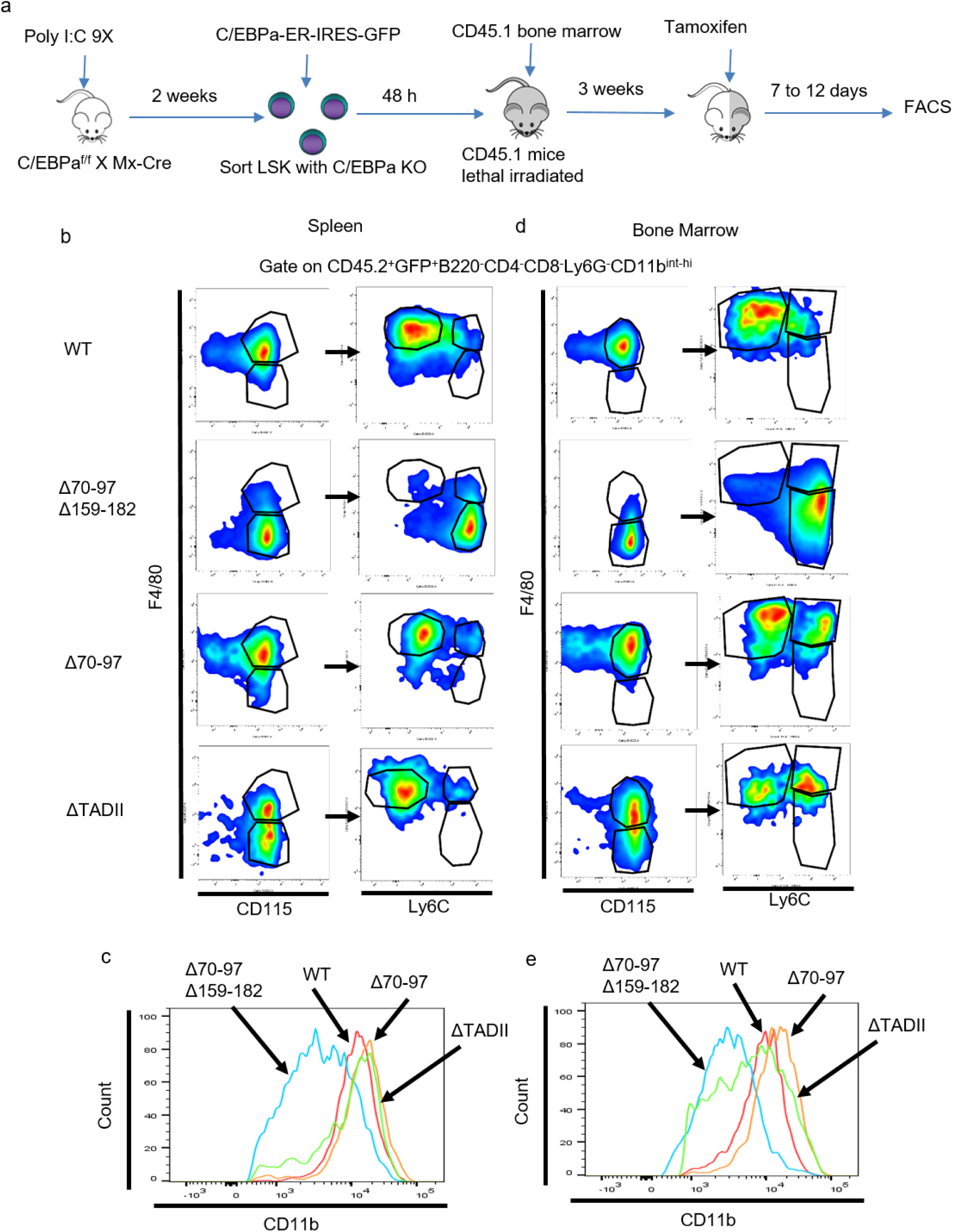
C/EBPα RNA binding affinity mainly affect macrophage population *in vivo*. (a) Experiment procedure of mouse bone marrow transplantation experiment. (b, c) Decrease of macrophage population F4/80^+^CD115^+^Ly6C^low^ (b) and CD11b level (c) upon double RNA binding domain deletion but not upon single deletion in spleen. (d, e) Decrease of macrophage population F4/80^+^CD115^+^Ly6C^low^ (d) and CD11b level (e) upon double RNA binding domain deletion but not upon single deletion in bone marrow.

We next investigated the macrophage differentiation among wild type and deletion mutations. Interestingly, the macrophage population (CD11b^high^CD115^+^F4/80^high^Ly6C^low^) rescued by C/EBPα wild type was disappeared and replaced by a monocyte population (CD11b^intermediate^CD115^+^F4/80^-^Ly6C^high^) in C/EBPα RNA binding domain deletions (Δ70-97+Δ159-182) in both spleen (Fig. 4b-c) and bone marrow (Fig. 4d-e), suggesting that C/EBPα RNA binding activity may essential for monocyte to macrophage differentiation. Furthermore, the C/EBPα single deletion mutations (Δ70-97 or ΔTADII), which were capable of binding to RNA, were able to rescue macrophage population similar to C/EBPα wild type, suggesting that the regulation of macrophage differentiation is specific to RNA binding activity.

To further confirm, we next checked the expression of different macrophage related genes by qRT-PCR (Fig. 5) using sorted GFP positive spleen and bone marrow cell from bone marrow transplanted mice. We observed decrease of PU.1 expression, a transcription factor regulates macrophage differentiation (Dahl and Simon, 2003; Nagamura-Inoue et al., 2001), upon deletion of RNA binding domains (Fig. 5a). Besides, the macrophage regulated gene CD163 (Fig. 5b) and CD206 (Fig. 5c) (Murray et al., 2014) were also decreased upon RNA binding domains deletion. Furthermore, genes like iNOS, IL1β, TNFα, IL10, Arg-1 and CCL-2 which are important for macrophage function (Gordon and Martinez, 2010; Martinez et al., 2009; Sica and Mantovani, 2012) were also decreased upon RNA binding domains deletion (Fig. 5d). Taken together the disappearing of macrophage like cell when depleted RNA binding domains in giemsa staining of in vitro cytokine (MCSF/IL3) assay (Fig. 5e), and the decrease in CD11b (Fig. 5f) and F4/80 (Fig. 5g) in same cells. These results suggest that C/EBPα without RNA binding capability failed to induce functional macrophages.

**Figure. 5.**
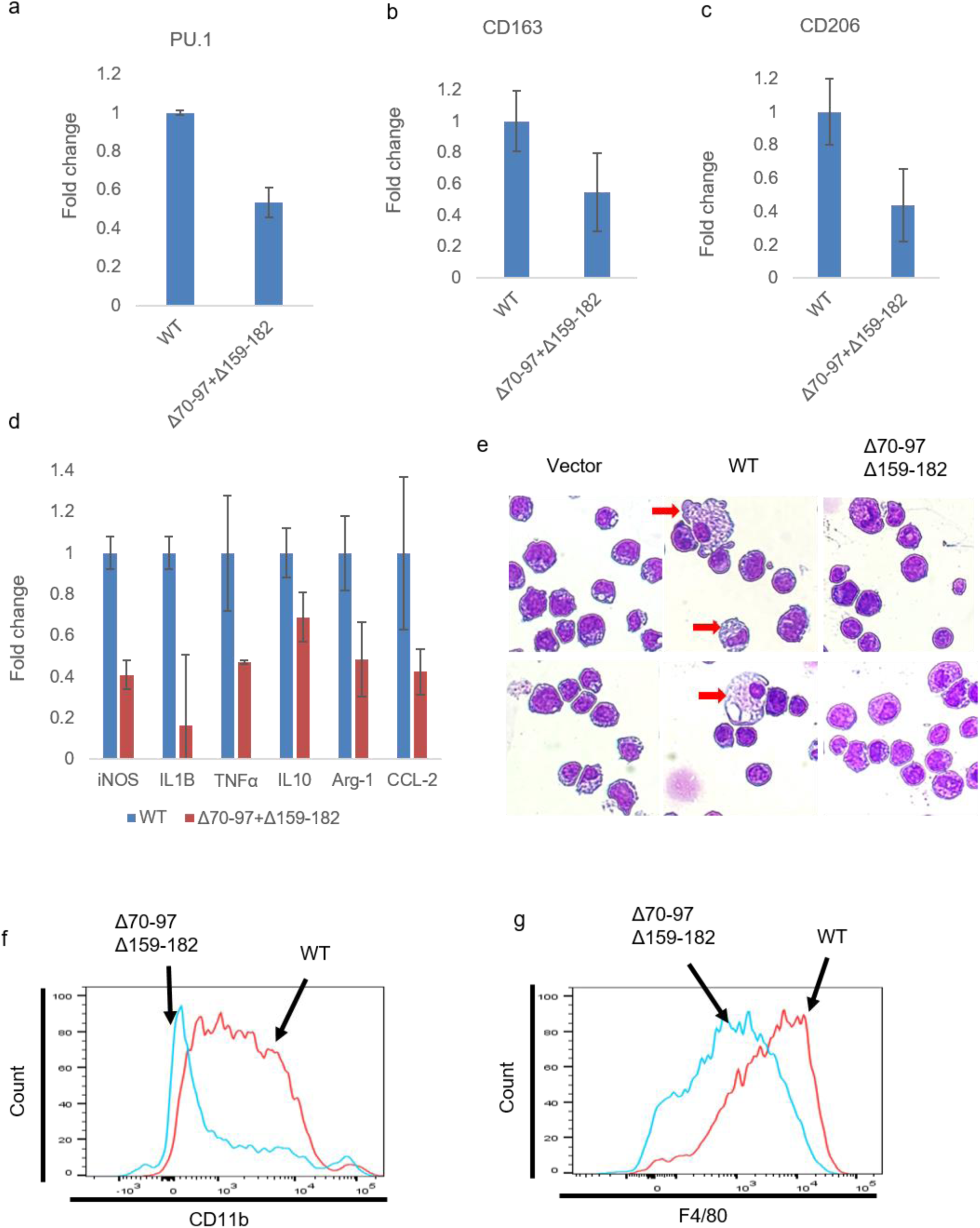
C/EBPα RNA binding activity regulates expression of macrophage related gene and promotes macrophage differentiation *in vitro*. (a-c) qRT-PCR analysis of GFP positive transplanted mouse primary spleen and bone marrow cells shows decrease expression of macrophage markers upon RNA binding domains deletion. PU.1 (a), CD163 (b), CD206 (c). (d) qRT-PCR analysis of GFP positive transplanted mouse primary spleen and bone marrow cells shows decrease expression of macrophage functional related gene upon RNA binding domains deletion. (e) Giemsa staining shows C/EBPα failed to induce macrophage like cells as C/EBPα did. (f, g) FACS analysis shows decrease of CD11b level (f) and F4/80 level (g) upon RNA binding domains deletion.

### C/EBPα RNA binding activity facilitates C/EBPα recruitment to PU.1 enhancer

PU.1 is essential for monocyte/macrophage differentiation and is regulated by C/EBPα through interacting with PU.1 enhancer (Yeamans et al., 2007), however how C/EBPα locate on specific site of PU.1 chromatin region is still remain unclear. Mouse primary GFP positive C/EBPα transplanted cells were sorted and ChIP-qRT-PCR was performed. Interestingly, enrichment of C/EBPα on PU.1 enhancer sites decreased upon RNA binding domain deletions (Fig. 6a). However, C/EBPα enrichment on C/EBPE and GCSFR sites, which are not C/EBPα RNA binding targets, are remain unchanged (Fig. 6a). Additionally, C/EBPα enrichment on PU.1 enhancer decreased upon overexpressing of C/EBPα double RNA binding domain deletion mutation in HL60 cells (fig. S7a-b).

**Figure. 6.**
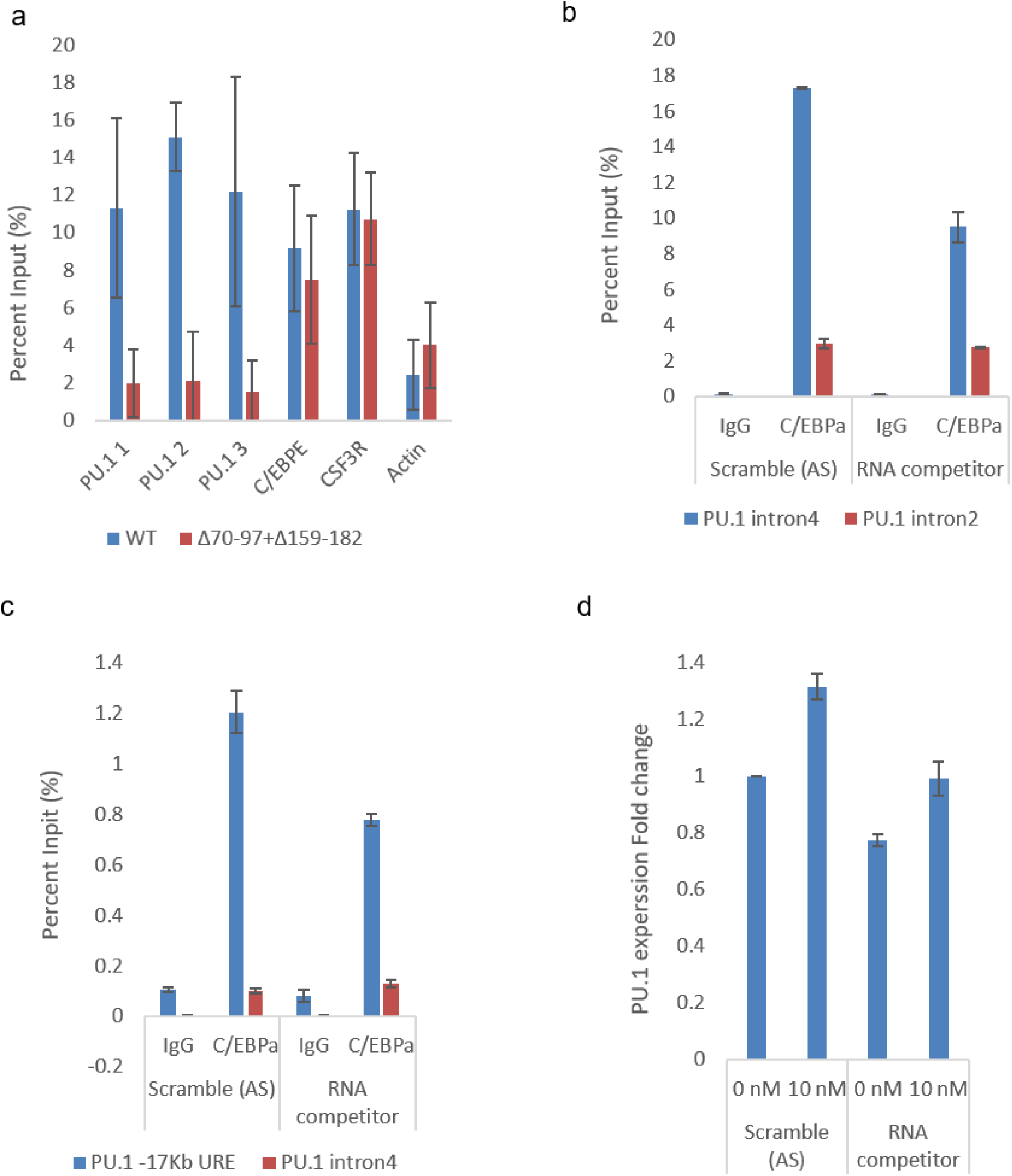
C/EBPα RNA binding activity promotes its chromatin recruitment. (a) ChIP-qRT-PCR analysis of GFP positive transplanted mouse primary spleen and bone marrow cells shows decreased enrichment of C/EBPα to PU.1 enhancer site but not C/EBPE and GCSFR (CSF3R) sites upon RNA binding domains deletion. (b) C/EBPα binding to PU.1 intron 4 decreased upon RNA competitor treatment. (c) C/EBPα recruitment to PU.1 enhancer site decreased upon RNA competitor treatment. (d) RNA competitor downregulates PU.1 expression level and inhibits PU.1 induction by PMA.

Besides, we also generated an in vitro transcribed RNA competitor and transfected in HL60 cells. Upon RNA competitor treatment, C/EBPα RNA binding on PU.1 intron 4 decreased (Fig. 6b), and correspondingly C/EBPα enrichment on PU.1 enhancer site decreased as well (Fig. 6c). However, there is no change on negative binding site in both RIP-qRT-PCR (Fig. 6b) and ChIP-qRT-PCR (Fig. 6c), suggesting that the change on binding site is specific. Correspondingly, PU.1 expression was also decreased upon RNA competitor treatment (Fig. 6d). Furthermore, we performed Phorbol 12-myristate 13-acetate (PMA) treatment assay in the context of with or without RNA competitor. PMA induced C/EBPα binding on PU.1 intron 4 RNA (fig. S7c) and PU.1 enhancer DNA (fig. S7d), and induced PU.1 expression (Fig. 6d). Interestingly, RNA competitor treatment blocked PMA induction on C/EBPα RNA binding (fig. S7c) and DNA binding (fig. S7d), and blocked PMA induction on PU.1 expression (Fig. 6d). The results suggested that C/EBPα RNA binding activity regulates the recruitment of C/EBPα to specific genome locus.

## Discussion

### Role of C/EBPα RNA binding activity in transcription regulation underlying macrophage differentiation

C/EBPα is known as a master transcription factor in neutrophil differentiation (Zhang et al., 1997; Zhang et al., 2004). Overexpressing C/EBPα-ER or C/EBPα bZIP domain mutation in cell line mainly affect neutrophil differentiation (D’Alo et al., 2003). However, C/EBPα express at a high level in monocyte/macrophage cells, indicating a potential role of C/EBPα in macrophage differentiation. Recent in vitro cell line studies have shown that C/EBPα regulates PU.1 expression through directly binds to PU.1 enhancer which may be important for monocyte differentiation (Yeamans et al., 2007). Additionally, overexpressing C/EBPα in *in vitro* cultured mouse primary B cell induces trans-differentiation into macrophage cells (Di Stefano et al., 2014; Hanna et al., 2008). Nonetheless, it remains poorly understood how C/EBPα promotes macrophage differentiation. Therefore it is intriguing to discover that the RNA binding activity of C/EBPα promotes macrophage differentiation which is mechanically and functionally different from its DNA binding activity. In our study, we identified two novel RNA binding domain located on N-terminal region of C/EBPα. Additionally, mouse in vivo bone marrow transplantation data showed that disrupt RNA binding activity mainly reduce macrophage population whereas has minor effect on neutrophil differentiation. Our findings provide a mechanistic explanation for the high level expression of C/EBPα in macrophage cells. Importantly, RNA binding function may also apply to other transcription factor, such as RUNX1 (Trinh et al., 2021) or PU.1, which may also play an important role in hematopoiesis. Our study extends our knowledge regarding the mechanically and functionally capability of transcription factor.

### Transcription factor RNA binding activity facilitates its recruitment to specific genome locus

C/EBPα mainly interacts with enhancer and promoter DNA through bZIP domain for transcription regulation (Pabst et al., 2001; Yeamans et al., 2007). Our study shows the importance of RNA binding activity of C/EBPα as another key determination in its transcription regulation. Emerging evidence has revealed the RNA binding capability of traditional DNA binding protein with various functions (Di Ruscio et al., 2013; Holmes et al., 2020; Nikolaev et al., 2010; Sigova et al., 2015; Wai et al., 2016). Genome-wide ChIP-seq and Gro-seq analysis revealed transcription factor YY1 interacts with RNA and DNA on same transcription element, indicating a potential role of RNA binding activity in transcription factor chromatin recruitment. Employing unbiased genome-wide chromatin RIP sequencing, we uncovered that C/EBPα mainly bond to intron of RNA transcript. ChIP-qPCR analysis revealed that disrupting RNA binding activity of C/EBPα through either deletion mutations or RNA competitor reduced C/EBPα enrichment on PU.1 enhancer. Correspondingly, PU.1 expression level decreased upon disrupting C/EBPα RNA binding activity. Our study provide a novel mechanistic explanation on the dynamic recruitment of transcription factor to specific gene locus.

## Materials and methods

### Animals

All mice used in the study were house in a sterile barrier facility approved by the IUCAC at the Beth Israel Deaconess Medical Center. Mx1-Cre mediated site-specific recombination of the LoxP cassettes in C/EBPa conditional knockout C57 mice has been generated and described in detail previously (REF). Cre^+/+^LoxP^+/+^ mice was confirmed by analyzing genomic DNA isolated from tail. The primers used for genotyping were listed in Supplementary Table. 1. Phenotype of knockout of C/EBPa was confirmed by flow cytometry (Fig. S5).

B6.SJLPtprca Pepcb/BoyJ (Pep Boy) congenic recipients mice was breed in lab.

Only Male mice was used in the study.

### Cell line

HL60 cell was maintained in IMDM media (HyClone, Cat. No. SH30228.01) with 20% fetal bovine serum (Hyclone, Cat. No. SH30910.03). THP-1 cell was maintained in RPMI-1640 media (Corning, Cat. No. 10-040-CM) with 10% fetal bovine serum and 0.05 mM β-mercaptoethanol (Fisher scientific, Cat. No. BP176-100). K562 cell was maintained in RPMI-1640 media with 10% fetal bovine serum. 293T-ecotropic and 293T-Amphotropic were maintained in DMEM high glucose media (Corning, Cat. No. 10-013-CV) with 10% fetal bovine serum. All cells were incubated at 37 degree Celsius and 5% CO_2_.

### Chromatin RNA immunoprecipitation (RIP)

50 million HL60 cells or THP-1 cells were suspended in 25 ml RPMI-1640 media without fetal bovine serum. 1.7 ml 16% formaldehyde (Thermo scientific, #28908) was added into cell solution (final concentration 1% formaldehyde) and shake at room temperature for 10 min. Glycine (Fisher scientific, Cat. No. G48-212) was then added to a final concentration of 1.5 M to stop the crosslinking reaction. Crosslinked cells were washed twitch with phosphate buffered saline (PBS) contains 5% vanadyl complex (New England Biolabs, Cat. No. S1492S) and 1 mM Phenylmethanesulfonyl fluoride (Sigma-Aldrich, Cat. No. P7626). After washing, cells were lysed in low salt lysis buffer [10 mM Tris-HCl pH7.4 (Thermo Fisher scientific, Cat. No. 17926), 10 mM NaCl (Sigma-Aldrich, Cat. No. S7653), 0.5% NP-40 (Sigma-Aldrich, Cat. No. I8896), 1X protease inhibitor (Roche, Cat. No. 11873580001), 1X Placental Rnase inhibitor (New England Biolabs. Cat. No. M0307L)] and dounce homogenized to isolate pure nuclei. Isolated nuclei were then lysed in high salt lysis buffer [50 mM HEPES (Fisher scientific, Cat. No. BP310-1), 1 mM MgCl_2_ (Sigma-Aldrich, Cat. No. M8266S), 0.1 mM CaCl_2_ (Sigma-Aldrich, Cat. No. 223506), 150 mM NaCl, 1% Triton-X100 (Sigma-Aldrich, Cat. No. T9284), 1% Sodium deoxycholate (Sigma-Aldrich, Cat. No. D6750), 1% Sodium dodecyl sulfate (SDS, Sigma-Aldrich, Cat. No. L3771), 1X protease inhibitor, 1X Placental Rnase inhibitor] and sonicate using Bioruptor 300 (Diagenode) at high intensity for 60 cycles with 15 seconds ON and 45 seconds OFF each cycle. Spin down and collected the supernatant, 5 µg antibody (Table. S3) and 50 µl protein A magnetic beads (New England Biolabs, Cat. No. S1425S) were added to the supernatant and incubate overnight at 4 degree Celsius. Beads was washed 6 times with washing buffer (150 mM NaCl, 10 mM Tris-HCl pH 7.4, 1% Triton-X100, 0.5% NP-40, 1X protease inhibitor, 1X Placental Rnase inhibitor), and then elute and reverse crosslink by incubate overnight at 60 degree Celsius in elution buffer [10 mM Tris-HCl pH 7.4, 10 mM EDTA (Sigma-Aldrich, Cat. No. E5134), 1% SDS, 200 mM NaCl, 200 µg protease K (Roche, Cat. No. 03115828001)]. Remove beads and collect supernatant, RNA was first isolated using phenol pH4 (Sigma-Aldrich, Cat. No. P4682) and ethanol. Isolated RNA was treatment with DNase I (Roche, Cat. No. 04716728001) at 37 degree Celsius for 30 minutes to remove any contaminate DNA. The RNA was purified again using phenol pH4 and ethanol. GlycoBlue (Invitrogen, Cat. No. AM9516) was added to help precipitating RNA.

### UV-crosslinked chromatin RNA immunoprecipitation (CLIP)

50 million HL60 or THP-1 cells were washed once with PBS and reseed into 15 cm culture plate (Falcon Corning, Cat. No. 353025). Plates were placed on ice and removed lid for UV crosslink at 150 J/cm^2^ using UV stratalinker 2400 (Agilent). After crosslink, cells were lysed in low salt lysis buffer (10 mM Tris-HCl pH7.4, 10 mM NaCl, 0.5% NP-40, 1X protease inhibitor, 1X Placental Rnase inhibitor) and dounce homogenized to isolate pure nuclei. Isolated nuclei were then lysed in high salt lysis buffer (50 mM HEPES, 1 mM MgCl_2_, 0.1 mM CaCl_2_, 150 mM NaCl, 1% Triton-X100, 1% Sodium deoxycholate, 1% SDS, 1X protease inhibitor, 1X Placental Rnase inhibitor) and sonicate using Bioruptor 300 (Diagenode) at high intensity for 60 cycles with 15 seconds ON and 45 seconds OFF each cycle. Spin down and collected the supernatant, 5 µg antibody (Table. S3) and 50 µl protein A magnetic beads were added to the supernatant and incubate overnight at 4 degree Celsius. Beads was washed 6 times with washing buffer (150 mM NaCl, 10 mM Tris-HCl pH 7.4, 1% Triton-X100, 0.5% NP-40, 1X protease inhibitor, 1X Placental Rnase inhibitor). Washed beads were then treated with Micrococcal nucleases (New England Biolab, Cat. No. M0247S) at 37 degree Celsius for 5 minutes. Reaction were stopped by stopping buffer [10 mM Tris-HCl pH 7.5, 50 mM NaCl, 1 mM EDTA, 50% Glycerol (Invitrogen, Cat. No. 15514-029)]. ^32^P were then labeled to 5’ end of digested RNA by T4 Polynucleotide Kinase (PNK) (New England Biolabs, Cat. No. M0201S). After labelling, beads were eluted in Nupage loading buffer (ThermoFisher Scientific, Cat. No. NP0007) at 70 degree Celsius for 10 minutes and directly run on SDS-PAGE gel. After running, protein-RNA complexes were transferred to nitrocellulose membrane (Bio-Rad, Cat. No. 1620112) for developing. Based on the developing results, cut out the membrane contains protein-RNA complex and incubate with protease K at 60 degree Celsius for 1 hour. Collect the supernatant and RNA was isolated using phenol pH4 and ethanol. GlycoBlue (ThermoFisher Scientific, Cat. No. AM9515) was added to help precipitating RNA. RNA obtained from CLIP was used in qRT-PCR not in RNA sequencing.

### Chromatin immunoprecipitation (ChIP)

Three to five million cells were suspended in 15 ml serum free RPMI-1640 media. 1 ml 16% formaldehyde was added to cells and incubate 10 minutes at room temperature. Crosslinked cells were washed twitch with phosphate buffered saline and lysed in SDS lysis buffer (50 mM Tris-HCl pH 8.1, 10 mM EDTA, 1% SDS). The cells were then sonicate using Bioruptor 300 at high intensity for 60 cycles with 15 seconds ON and 45 seconds OFF each cycle. Spin down and collected supernatant, the supernatant was then diluted 10 times using ChIP dilution buffer (20 mM Tris-HCl pH 8.1, 150 mM NaCl, 1 mM EDTA, 1% Triton-X100). 5 µg antibody (Table. S3) and 50 µl protein A magnetic beads were added to the solution and incubate overnight at 4 degree Celsius. Beads were washed once with low salt wash buffer (20 mM Tris-HCl pH 8.1, 150 mM NaCl, 2 mM EDTA, 1% Triton-X100, 0.1% SDS), once with high salt wash buffer (20 mM Tris-HCl pH 8.1, 500 mM NaCl, 2 mM EDTA, 1% Triton-X100, 0.1% SDS), once with LiCl wash buffer [10 mM Tris-HCl pH 8.1, 1 mM EDTA, 1% Sodium deoxycholate, 1% NP-40, 250 mM lithium chloride (Sigma-Aldrich, Cat. No. L9650)] and twice with TE buffer (10 mM Tris-HCl pH 8.1, 1 mM EDTA). Beads were then eluted and reverse crosslinked by incubate overnight at 60 degree Celsius in elution buffer (10 mM Tris-HCl pH 7.4, 10 mM EDTA, 1% SDS, 200 mM NaCl, 200 µg protease K). Beads were removed and DNA was isolated using PCR purification kit (Qiagen, Cat. No. 28104).

### Real time quantitative polymerase chain reaction (RT-qPCR)

RNA obtained from RIP or CLIP, or mRNA isolated from cell lines or primary mouse cells were proceeded to RT-qPCR using iTaq universal SYBR green one step kit (Bio-Rad, Cat. No. 1725150). DNA obtained from ChIP was proceed to RT-qPCR using iQ SYBR green supermix (Bio-Rad, Cat. No. 1708880). Rotor-Gene 6000 platform (Corbett, US) was used for data acquisition and analysis. Primers were listed in Supplementary Table. 1.

### RNA sequencing (RNAseq)

RNA obtained from RIP was proceed for RNAseq. RNAseq library was prepared using TruSeq stranded total RNA sample preparation kits (illumine, Cat. No. 20020597) following manufactory protocol. The library was validated on 2100 Bioanalyzer instrument (Agilent, Santa Clara, CA) and sequenced using Nextseq500.

### Protein purification

His-tagged or His-sumo-tagged protein was purified using NiNTA beads (Anatrace, Cat. No. SUPER-NINTA25). Briefly, Bacteria transformed with expression constructs were cultured at 37 degree Celsius till OD600 reached 0.8 to 1.0. 1 mM IPTG (Sigma-Aldrich, Cat. No. I6758) was added to bacteria culture and incubate overnight at room temperature. Next day, bacteria cells were spin down and suspended in buffer H (50 mM Tris-HCl pH 8.0, 150 mM NaCl, 10% Glycerol) containing 1 mg/ml lysozyme (Bio Basic, Cat. No. LDB0308-5) and 0.1 % TritonX-100. Cells were incubated on ice for 30 minutes and sonicate using sonic dismembrator model 500 (fisher scientific) for 20 minutes (30s ON, 30s OFF, 30% amplitude). The solution became transparent after sonication, then spin down at 13,000 rpm for 15 minutes. Collect supernatant and added 1 ml NiNTA beads (50% slurry, washed 3 times using buffer H). Incubated at 4 degree Celsius for 2 hours. Beads were then washed twice with buffer H containing 20 mM imidazole (Sigma-Aldrich, Cat. No. I2399) and twice with buffer H containing 40 mM imidazole. Washed beads were eluted using buffer H containing 300 mM imidazole. Eluted protein was dialyzed into different buffers based on following experiments using snakeskin dialysis tubing 3.5K MWCO 16 mm (Thermo scientific, Cat. No. 88242).

For RNA EMSA, protein was dialyzed into REMSA buffer [10mM HEPES pH 7.3 (Sigma-Aldrich, Cat. No. H3375), 20 mM KCl (Fisher Scientific, Cat. No. P217), 1 mM MgCl_2_, 1 mM DTT (GoldBio, Cat. No. DTT10), 1 mM EDTA, 10% Glycerol].

For ITC, protein was dialyzed into ITC buffer (20 mM HEPES pH 7.5, 150 mM NaCl, 1 mM MgCl_2_, 2 mM DTT).

For NMR, protein was dialyzed into NMR buffer [30 mM phosphate buffer pH 7.4 (22.6 mM Na_2_HPO_4_ (Sigma-Aldrich, Cat. No. S9763) and 7.4 mM NaH_2_PO_4_ (Sigma-Aldrich, Cat. No. S0751), 150 mM NaCl, 1 mM MgCl_2_, 2 mM DTT].

### Transfection and retrovirus stock preparation

Retrovirus plasmids were transfected into 293T-Eco cells using Lipofectamine 2000 (Thermo scientific, Cat. No. 11668019). 48 hours post transfection, supernatant was harvested and filtered by 0.45 µm hydrophilic PVDF filter (Foxx life science, Cat. No. 378-3415-OEM). Retro-X concentrator (Takara, Cat. No. 631456) was added to filtered supernatant at a ratio 1:3 and incubated overnight at 4 degree Celsius. After incubation, spindown and suspended the virus pallet in 500 µl StemSpan SFEM serum free media (stemcell technologies, Cat. No. 09650). Make a small aliquot for virus titration using qPCR titration kit (abm, Cat. No. G949). The remaining virus was put in −80 degree celsius freezer.

### Retrovirus transduction and bone marrow transplantation assay

C/EBPα^f/f^ Mx1-Cre^+^ mice were intraperitoneal injected 30 µg Poly I:C (Sigma-Aldrich, Cat. No. P0913) per gram of mouse weight every alternative day for 18 days to induce C/EBPα conditional knockout. Two tibia, two femur and two pelvis were harvested from each knockout mice. Bone marrow cells were isolated from each bones and proceed to lineage depletion using direct lineage depletion kit (Milenyi Biotec, Cat. No. 130-110-470). Cells were then stained with antibody (Supplementary Table. 2) and LSK cells were isolated through cell sorting using FACSAria (BD). Sorted LSK cells were cultured in StemSpan SFEM serum free media (stemcell technologies) contains 40 ng/ml mSCF (Biolegend, Cat. No. 579702), 40 ng/ml mTPO (Biolegend, Cat. No. 593302) and 10 ng/ml IL3 (Biolegend, Cat. No. 575502) for virus transduction. 200 MOI retrovirus and 5 µg/cm^2^ RetroNectin reagent (Takara, Cat. No. T100A) were used in the transduction following the manufactory protocol. 48 hours post transduction, 1 × 10^5^ LSK cells per mouse were injected into lethally irradiated (1000 cGy) B6.SJLPtprca Pepcb/BoyJ (Pep Boy) congenic recipients along with 1 × 10^5^ lineage depleted Pep boy bone marrow cells for radioprotection.

### Flow cytometry (FACS)

Single-cell suspension isolated from various organs were analyzed by FACS. Antibody used were listed in Supplementary Table. 2. Stained cells were analyzed using LSRII flow cytometer (BD Biosciences, San Jose, CA), LSR fortessa flow cytometer (BD) or CytoFLEX LX flow cytometer (BECKMAN COULTER life sciences, Indianapolis, IN). DIVE software (BD), CytoExpert (Beckman) and FlowJo V10 (Tree Star) were used for data acquisition and analysis.

### Primary cell sorting

Single-cell suspension received from primary cell isolation were proceeded to cell sorting. Antibody used were listed in Supplementary Table. 2. Stained cell were suspended at a concentration of 20 million cell per milliliter in phosphate-buffered saline (PBS) plus 0.5% fetal bovine serum (FBS). LSK cell or GFP positive CD45.2 cell were obtained using FACSAria (BD).

### Western blot

Cells were lysed using 1X RIPA buffer (ThermoFisher Scientific, Cat. No. 89900) with 1X protease inhibitor cocktail follow by 3 times sonication (Sonic Dismembrator Model 100, Fisher Scientific, Waltham, MA) with 10 seconds each time at intensity level 3. Protein was quantified using BCA protein assay kit (Thermo scientific, Cat. No. 23227) following the manufactory protocol. Protein lysates were separated on 10% self-made Bis-tris gel and transferred to nitrocellulose membranes. The membrane was blocked by 5% Bovine Serum Albumin (bioworld, Cat. No. 22070008) in 1X TBST for 1 hour at room temperature and incubated with primary antibody (Supplementary Table. 2) overnight at 4 degree Celsius. The membrane was then incubated with Secondary antibody (Supplementary Table. 2) 1 hour at room temperature, developed using reagent (ThermoFisher Scientific, Cat. No. 34580), and observed on Amersham Imager 600 (GE Healthcare).

### RNA electrophoretic mobility shift assay

C/EBPα full length and mutations were purified from bacteria and dialyzed in REMSA buffer. 19-21 bp RNA probe were obtained from PU.1 intron 4 (PU.1i4): CCCCCACCCAGCCAGGCCAAC and CTBP1 intron 2 (CTBP1i2): UUCCCACUCACCCAGCACCC. A negative binding probe was also obtained from CTBP1 intron 2 (CTBP1i2N): GGGACCCGGUUCCCACUCACCCAGC. RNA probes were purchased from Sigma-Aldrich and proceed to 5’end ^32^P labelling using PNK. After labelling, labelled probe was purified by native TBE-PAGE gel and illustra G-25 columns (GE Healthcare, Cat. No. 27532501). Equal intensity of different RNA probes were mixed with equal molar of different C/EBPα proteins individually in EMSA buffer B1 (Active Motif, Cat. No. 37480), stabilizing solution D (Active Motif, Cat. No. 37488) and buffer C1 (Active Motif, Cat. No. 37484). After incubating on ice for 10 minutes, the mixture was running on 4% native TBE-PAGE gel. After running, gel was fixed by fixing buffer [20% Methanol (Pharmco, Cat. No. 339000000), 10% Acetic acid (Sigma-Aldrich, Cat. No. 695092)] and dried on Model 583 gel dryer (Bio-Rad). The dried gel was then developed directly using X-ray film (MTC bio, Cat. No. A8815). For overnight exposure, the cassette was put in −80 degree Celsius freezer.

### In vitro transcription and electroporation

Linearized DNA plasmids were proceeded to in vitro transcription using MAXIscript SP6 transcription kit (ThermoFisher Scientific, Cat. No. AM1308). After reaction, DNAse I was added to remove template DNA. RNA was isolated by phenol pH4 and ethanol. The quality of RNA was examined by running 2% agarose gel. The sequence of RNA are as follow: RNA competitor: 5’-ACC CTG GCC CCG ACC CTG GCC GTG TCT GTG CCC TGC CCC CAC CCA GCC AGG CCA ACT GAC CCA GCA GCC A-3’; Scramble (anti-sense): 5’-TGG CTG CTG GGT CAG TTG GCC TGG CTG GGT GGG GGC AGG GCA CAG ACA CGG CCA GGG TCG GGG CCA GGG T-3’.

Electroporation was performed using nucleofector kit V (Lonza, Cat. No. VCA-1003) and Amaxa nucleofector II electropotator (Lonza) following the manufactory protocol. 2 million HL60 cells and 5 µg in vitro transcribed RNA were used. 24 hours post electroporation, cells were harvested for ChIP, RIP and qRT-PCR.

